# Pharmacophore-guided discovery of CDC25 inhibitors causing cell cycle arrest and cell death

**DOI:** 10.1101/309914

**Authors:** Zeynep Kabakci, Simon Käppeli, Giorgio Cozza, Claudio Cantù, Christiane König, Janine Toggweiler, Christian Gentili, Giovanni Ribaudo, Giuseppe Zagotto, Konrad Basler, Lorenzo A. Pinna, Stefano Ferrari

**Author notes:** These authors contributed equally to the study (Zeynep Kabakci, Simon Käppeli).

## Abstract

CDC25 phosphatases have a key role in cell cycle transitions and are important targets for cancer therapy. Here, we set out to discover novel CDC25 inhibitors. Using a combination of computational approaches we defined a minimal common pharmacophore in established CDC25 inhibitors and performed a virtual screening of a proprietary library. Taking advantage of the availability of crystal structures for CDC25A and CDC25B and using a molecular docking strategy, we carried out hit expansion/optimization. Enzymatic assays revealed that naphthoquinone scaffolds were the most promising CDC25 inhibitors among selected hits. At the molecular level, the compounds acted through a mixed-type mechanism of inhibition of phosphatase activity, involving reversible oxidation of cysteine residues. In 2D cell cultures, the compounds caused arrest of the cell cycle at the G1/S or at the G2/M transition. Mitotic markers analysis and time-lapse microscopy confirmed that CDK1 activity was impaired and that mitotic arrest was followed by death. Finally, studies on 3D organoids derived from intestinal crypt stem cells of *Apc/K-Ras* mice revealed that the compounds caused arrest of proliferation.

## INTRODUCTION

The Cell Division Cycle 25 family encompasses three highly conserved members of dual specificity phosphatases that specifically target Cyclin-Dependent Kinases (CDKs), acting as dose-dependent inducers of cell cycle transitions (Dunphy & Kumagai, 1991, Russell & Nurse, 1986). CDC25A primarily activates CDK2/CycE and CDK2/CycA at the G1/S transition and in S-phase (Hoffmann, Draetta et al., 1994), though it also cooperates with CDC25B at the onset of mitosis (Lindqvist, Kallstrom et al., 2005). CDC25B initiates CDK1/CycB activation at centrosomes during the G2/M transition (Gabrielli, De Souza et al., 1996, Lindqvist et al., 2005) and CDC25C causes full activation of CDK1 at mitotic entry (Gabrielli, Clark et al., 1997).

Genetic studies showed that thermosensitive *cdc25* yeast mutants could be reversibly arrested in the cell cycle (Kovelman & Russell, 1996), providing the first demonstration of a regulatory role for CDC25. The mouse *Cdc25A* gene was shown to be the only family member endowed with an essential function during embryonic development (Lee, White et al., 2009).

Overexpression of CDC25, particularly CDC25A and CDC25B, has been observed in a variety of human cancers and correlates with poor clinical prognosis (Boutros, Lobjois et al., 2007). Interestingly, although CDC25A overexpression alone is insufficient to drive tumor initiation, *CDC25A* has a clear role as rate-limiting oncogene in transformation by *RAS* (Ray & Kiyokawa, 2008). Furthermore, point mutations in CDC25C have a critical role in the pathology of acute myelogenous leukemia (AML) (Yoshimi, Toya et al., 2014).

Since the identification of vitamin K as potent CDC25 inhibitor, compounds based on the structure of vitamin K, as well as on other structures have been developed as CDC25 inhibitors (Bana, Sibille et al., 2015, Brezak, Quaranta et al., 2004, Brun, Braud et al., 2005, Contour-Galcera, Lavergne et al., 2004, Kar, Lefterov et al., 2003, Lavecchia, Di Giovanni et al., 2012b, Pestell, Ducruet et al., 2000, Song, Lin et al., 2014). However, the compounds so far reported have failed to keep up with expectations, either due to rapid metabolism in tumor-bearing SCID mice (Guo, Parise et al., 2007) or for not completing clinical trials (Lavecchia, Di Giovanni et al., 2010), and have thus not attained approval.

In light of the recent discovery that CDC25 is the therapeutic target of choice in triple-negative breast cancers, namely those that are negative for estrogen-, progesterone- and HER2-receptor expression and that are unresponsive to standard therapy (Liu, Granieri et al., 2018), we set out to develop novel CDC25 inhibitors. To this end, we conducted a pharmacophore-guided drug discovery program that led to the identification of novel scaffolds of the naphthoquinone group displaying inhibition of CDC25 in enzymatic assays. In cultured cells, the most potent scaffolds induced inhibition of CDK1 activity and function, with block of mitotic transition followed by cell death. In mouse *Apc/K-Ras* mutant duodenal organoids, low doses of CDC25 inhibitors caused arrest of proliferation and expression of differentiation markers, whereas high doses induced cell death. The data reveal the potential of these novel compounds, used either as single agent or in combination with other drugs, to target tumors that depend on CDC25.

## RESULTS

### Pharmacophore-guided library screening and hit selection

To the end of retrieving novel CDC25 inhibitors from an *in silico* virtual library that was built from a proprietary database of synthetic molecules, we implemented a number of computational strategies (Fig. S1A), according to established protocols (Koch, Schuffenhauer et al., 2005). First, CDC25 inhibitors belonging to three classes - natural products, quinones and electrophiles (Lazo & Wipf, 2008) - were subjected to a linear fragmentation process (Schuffenhauer, Ertl et al., 2007) implemented in MOE Suite (G., 2010), in which input structures were split into small pieces by removing the least "scaffold-like" extremity until indivisible essential fragments were obtained. Next, the molecular entities returned by this process, ordered by increasing size, were used to build a series of pharmacophore models (Fig. S1B). The latter were optimized until the achievement of a final model, representative of the chemical features of scaffolds obtained from the fragmentation process. Finally, this model was used to examine a proprietary library through a pharmacophore-guided virtual screening process (MOE Suite). Compounds obtained from the first round of hit selection and belonging to different molecular families were tested at fixed concentration on recombinant CDC25A (Table S1). Reference compound in all tests was the established CDC25 inhibitor NSC-663284, a para-quinonoid derivative of vitamin K (Lazo, Aslan et al., 2001). The naphthoquinones UPD-140 (Fig. 1A, 2-(2’,4’-dihydroxyphenyl)-8-hydroxy-1,4-naphthoquinone) and UPD-176 (Fig. 1B, 5-hydroxy-2-(2,4-dihydroxyphenyl)naphthalene-1,4-dione) appeared to be the most effective inhibitors of CDC25 phosphatase activity. Based on the structure of UPD-140 and UPD-176, and exploiting the crystal structure of CDC25B (Reynolds, Yem et al., 1999), along with available homology models for CDC25A and CDC25C, we performed hit expansion/optimization through a molecular docking strategy (Fig. 1C). Identification of pockets and surface sites through the localization of regions of tight atomic packing suggested two close cavities suitable to accommodate the compounds. Both cavities are highly conserved in the three enzymes and one is superimposable with the phosphatase catalytic site (Table 1). Starting point for prioritization of scaffolds was the presence of a quinone moiety, which appeared to be a necessary condition for optimal anchoring of compounds in CDC25 catalytic pocket. Reassessment of the library, based on the structure of UPD-140 and UPD-176, followed by in *vitro* enzymatic assays revealed eight additional compounds as effective inhibitors of CDC25 phosphatase activity, all of them being 1,4-naphthoquinones with hydroxyl groups either in position 5 or 8 (Table S2).

**Figure 1.**
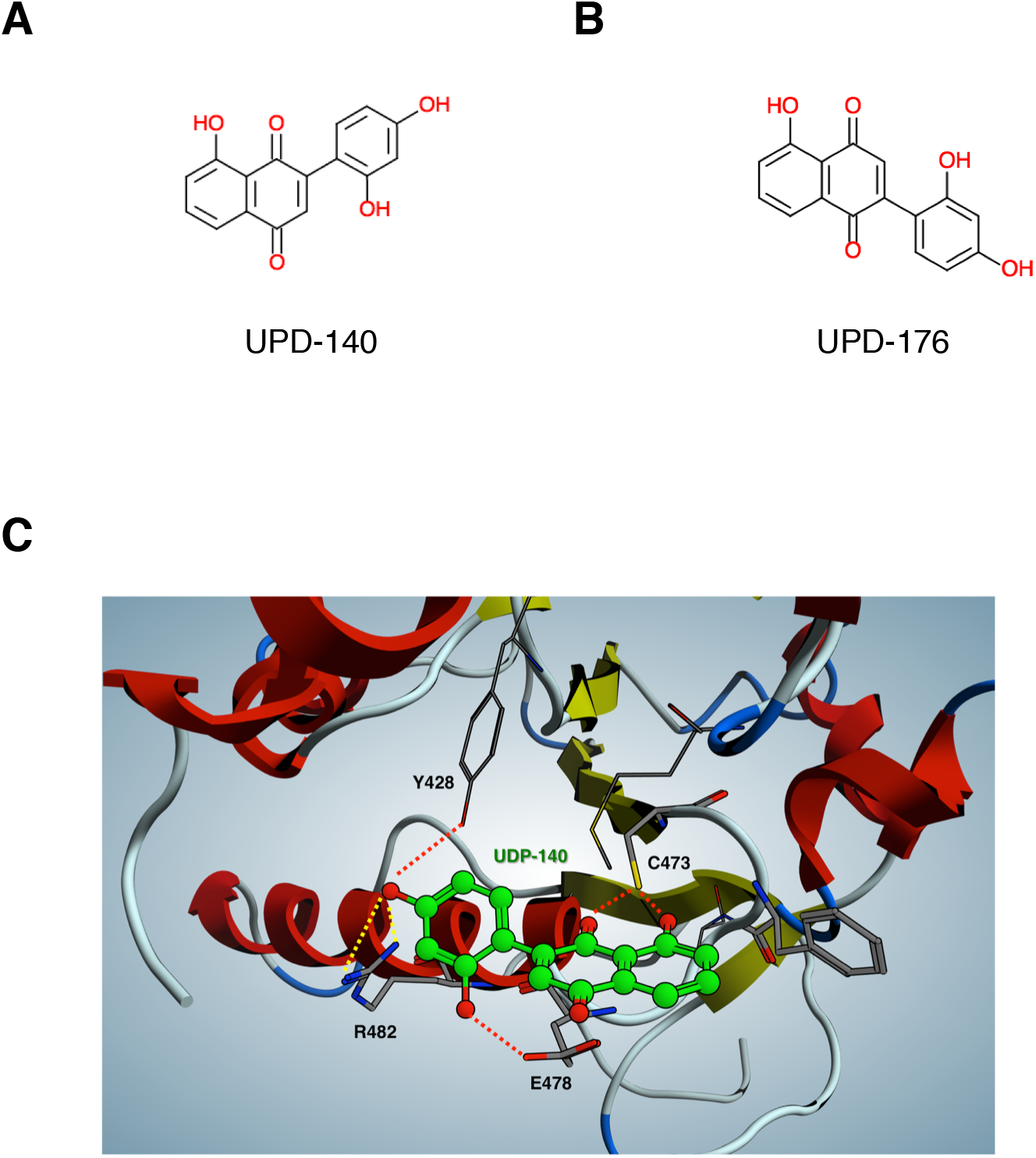
Structure and docking of CDC25 inhibitors. (A, B) Structure of 2-(2’,4’-dihydroxyphenyl)-8-hydroxy-1,4-naphthoquinone (UPD-140) and 5-hydroxy-2-(2,4-dihydroxyphenyl)naphthalene-1,4-dione (UPD-176). (C) Molecular docking of UPD-140 into CDC25B catalytic site.

**Table 1.** Residues lining the two cavities of CDC25 retrieved by the Site Finder approach. The catalytic cysteine in cavity 1 of each phosphatase is shown in italics.

| Cavity 1 |  |  | Cavity 2 |  |  |
| --- | --- | --- | --- | --- | --- |
| **CDC25A** | **CDC25B** | **CDC25C** | **CDC25A** | **CDC25B** | **CDC25C** |
| Tyr 387 | Tyr 428 | Tyr 332 | Asp 336 | Glu 377 | His 281 |
| *Cys 431* | *Cys 473* | *Cys 377* | Ile 338 | Ile 379 | Ile 283 |
| Phe 433 | Phe 475 | Phe 379 | Gly 339 | Gly 380 | Gly 284 |
| Ser 434 | Ser 476 | Ser 380 | Tyr 345 | Phe 386 | Cys 290 |
| Glu 436 | Glu 478 | Glu 382 | Asp 356 | Asp 397 | Asp 301 |
| Gly 438 | Gly 480 | Gly 384 | Leu 357 | Leu 398 | Leu 302 |
| Arg 440 | Arg 482 | Arg 386 | Lys 358 | Lys 399 | Lys 303 |
| Met 489 | Met 531 | Met 435 | Cys 442 | Cys 484 | Cys 388 |
|  |  |  | Arg 446 | Arg 488 | Arg 392 |
|  |  |  | Arg 450 | Arg 492 | Arg 396 |
|  |  |  | Glu 462 | Glu 504 | Glu 408 |
|  |  |  | Leu 463 | Met 505 | Leu 409 |

### Enzymatic characterization and profiling of CDC25 inhibitors

Selected naphthoquinone-based inhibitors were examined in dose-response assays on CDC25 phosphatases. Compounds UPD-786, UPD-793, UPD-795 and UPD-140 appeared to be the most potent inhibitors, displaying IC_50_ of 0.89 μM, 0.92 μM, 1.25 μM and 1.42 μM on CDC25A that are comparable to the IC_50_ of NSC-663284 (0.38 μM) (Fig. 2A and S2A) (Pu, Amoscato et al., 2002). A similar pattern of inhibition was observed on CDC25B and CDC25C, though with slightly higher IC_50_ values (Fig. 2A). Kinetic analysis conducted with UPD-795 revealed a mixed mechanism of enzyme inhibition, as indicated upon data analysis with the mixed-model inhibition equation (Copeland, 2005) (Table S3 and Fig. S2B), confirming the predictions of docking studies.

**Figure 2.**
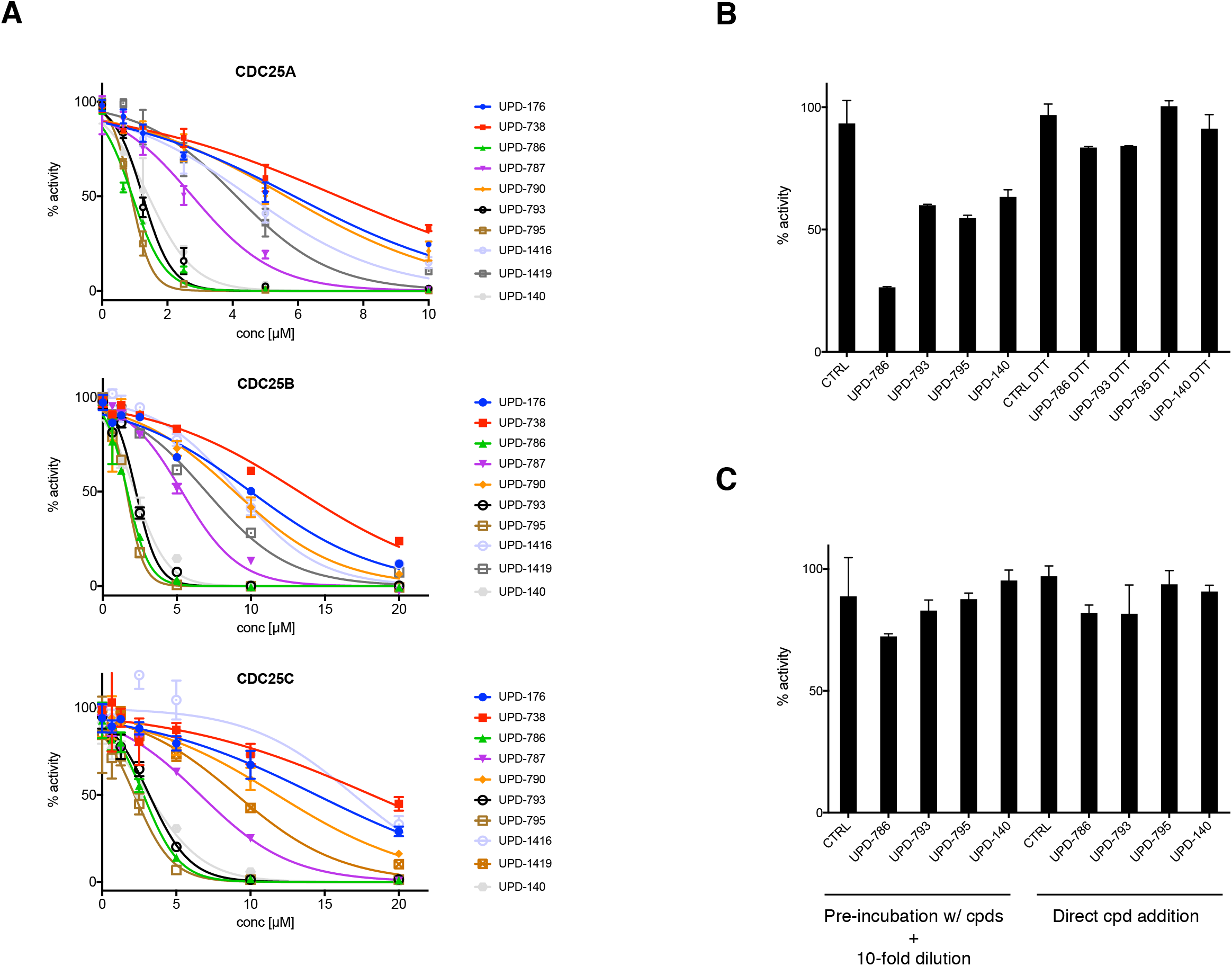
Enzymatic analysis of CDC25 inhibitors. (A) Dose-response studies with the indicated compounds on recombinant CDC25A, CDC25B and CDC25C. (B) CDC25A was treated in the presence or the absence of the reducing agent DTT (10 mM) prior to incubation with the compounds at a concentration proximal to their IC50 (1.25 μM). (C) CDC25A was treated with the indicated compounds at a concentration ~3x IC_50_ (5 μM), diluted and assayed for phosphatase activity at the remaining compound concentration (0.5 μM). As comparison, CDC25A was assayed upon direct addition of compounds at 0.5 μM concentration.

Considering that quinones are oxidizing agents, known for their ability to generate reactive oxygen species (ROS) in biological systems, and that the Cys residue in CDC25 active site is highly susceptible to oxidation (Sohn & Rudolph, 2003), we decided to assess a possible role for redox reactions in the inhibition of CDC25 by the most potent naphthoquinones described here. To this end, we treated CDC25A with an excess of the reducing agent DTT before addition of the compounds at a concentration proximal to their IC_50_. The data indicated that the presence of DTT in the reaction prevented inhibition of CDC25 (Fig. 2B).

Furthermore, since the quinone scaffolds so far described in the literature are in part reversible and in part irreversible CDC25 inhibitors, we decided to assess the mode of action of our most potent naphthoquinone inhibitors. To this end, we pre-incubated CDC25A with excess compound (5 μM, corresponding to ~3x IC_50_) and subsequently assayed for remaining phosphatase activity upon 10-fold dilution of the preincubation mix (hence at 0.5 μM compound). The data showed that the enzyme activity remaining upon dilution of the pre-incubation mix was comparable to the activity detected upon direct treatment of the phosphatase with 0.5 μM compounds, indicating a reversible mode of action (Fig. 2C).

To assess specificity, we profiled two of the most potent CDC25 inhibitors, UPD-795 and UPD-140, against a panel of protein phosphatases. The data revealed that, when tested at concentrations close to the IC_50_ for CDC25A, both compounds inhibited human PP5 activity to about 50% (Table S4), whereas none of the other phosphatases examined was affected.

### Cellular characterization of CDC25 inhibitors

To assess CDC25 inhibitors in cells we performed viability assays on HeLa cells upon treatment with increasing compound doses. The data revealed IC_50_ of 1.08 μM, 1.41 μM, 0.90 μM and 1.20 μM for UPD-176, UPD-790, UPD-787 and UPD-140 respectively (Fig. 3A), all values being lower than that of the reference compound NSC-663284 (IC50 = 2.57 μM) (Fig. S3A). Compounds UPD-738 and UPD-786 appeared to be slightly less potent than NSC-663284, though both displayed IC_50_ values <10 μM (4.40 μM and 6.50 μM, respectively) (Fig. S3B).

**Figure 3.**
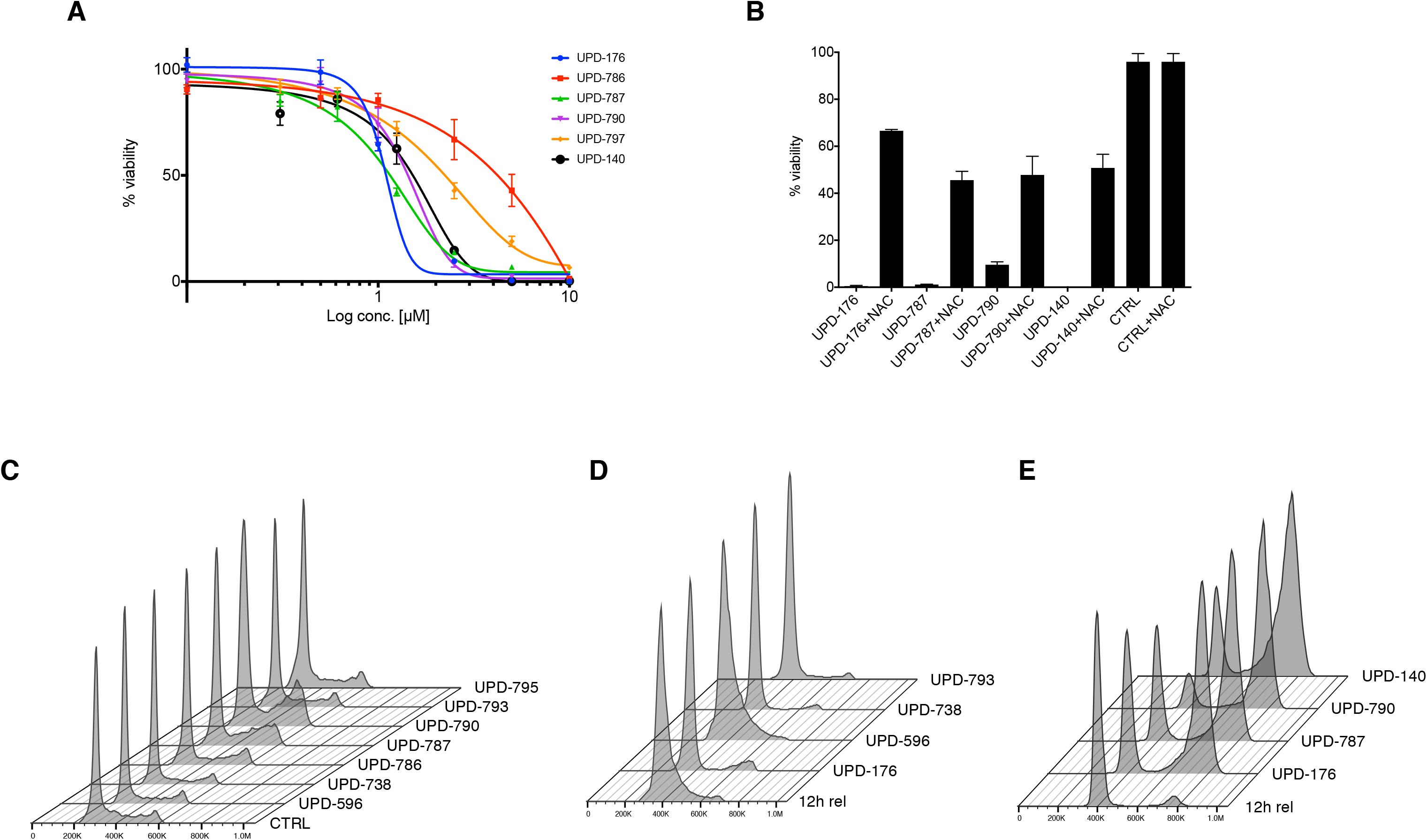
CDC25 inhibitors affect cell viability and the cell cycle. (A) HeLa cells were treated with increasing amounts of compounds and cell viability was determined. (B) Cells were treated in the presence or the absence of NAC before administration of compounds at doses ~3x IC_50_ (5 μM) and cell viability was determined. (C) Cells were treated with the indicated compounds (10 μM, 15h). DNA was stained with DAPI. (D) HeLa cells were synchronized at G2/M by treatment with Thymidine (1 mM, 15h), release in full medium followed by addition of the CDK1 inhibitor Ro-3306 (9 μM, 15h). Five hours upon release from Ro-3306, the indicated compounds were added (10 μM) and cells were harvested and analyzed 12h upon release from Ro-3306. (E) HeLa cells were synchronized at G1/S by 2x thymidine block-release, compounds (10 μM) were added at 5h and cells were analyzed at 12h from release.

To evaluate in a cellular context the oxidation-based mechanism of inhibition, we treated cells with the aminothiol N-Acetylcysteine, an antioxidant, prior to administration of the compounds. At compound doses exceeding 3x the IC_50_ (5 μM), cell viability was rescued to ~50% of controls in the presence of the reducing agent (Fig. 3B), hence confirming results obtained in biochemical assays.

To assess effects of the CDC25 inhibitors on cell cycle progression we administered compounds to HeLa cells and examined DNA content by flow cytometric analysis. In asynchronous HeLa cells UPD-738 and, to a lesser extent, UPD-786 and UPD-793 caused a decrease of the S-phase cell population, indicating that G1 was the point of action. On the other hand, UPD-787 and UPD-790 were most effective in causing accumulation of cells at G2/M (Fig. 3C). Similar results were obtained in U2OS cells (Fig. S4). Since these model systems can be conveniently synchronized at specific points in the cell cycle (Fig. S5A), HeLa cells were treated with UPD-176 or UPD-795 at the time of release from a double-thymidine block (early S-phase). Under these conditions, progression through S-phase was effectively held in check as compared to controls (Fig. S5B). To precisely assess effects of the compounds on the G1/S transition, HeLa cells were synchronized at G2/M, released to allow completion of mitosis and re-entry into the cell cycle, and treated in mid-G1 with UPD-176, UPD-596, UPD-738 or UPD-793. Analysis of cell cycle progression at 12h post-release, a time when control cells entered S-phase, showed that UPD-176, UPD-738 and UPD-793 effectively blocked cells in G1, whereas UPD-596 appeared to be ineffective in this respect (Fig. 3D). To assess the effect of compounds on the G2/M transition, double-thymidine synchronized HeLa cells were released and treated with UPD-176, UPD-787, UPD-790 or UPD-140 5h upon release, namely in late S-phase (Fig. S5A). Flow cytometric analysis of cell cycle progression at 12h post-release, when control cells largely moved to the next G1 phase, showed that all compounds caused accretion of the G2/M peak, with UPD-790 apparently being the most effective of all (Fig. 3E). To exclude the possibility that the observed cell cycle arrest would be secondary to genotoxic effects, we examined the biomarker γ-H2AX at the onset of mitosis in cells treated with the compounds. The data showed that, in comparison to a potent genotoxic agent, no damage to DNA occurred upon treatment with the naphthoquinone compounds (Fig. S6).

To visualize the execution of mitosis, we administered compounds to HeLa cells 5h upon release from a double-thymidine block, and monitored cells by time-lapse microscopy for the subsequent 14h. We observed that whereas control cells timely progressed through mitosis and moved to the next cycle, cells treated with UPD-787 (Fig. 4A) could not complete mitotic transition and died before reaching G1. A similar pattern was obtained with compounds UPD-790 and UPD-176 (Fig. S7 and data not shown). To better appreciate the response to inhibitors at the onset of mitosis, we administered low doses (5 μM) of UPD-787 or UPD-790 to synchronized HeLa Kyoto cells, which carry mCherry-H2B and GFP-a-tubulin. We observed a failed attempt to round up and execute mitosis, which was followed by membrane blebbing and death (Fig. S7 and movies M1-M3).

**Figure 4.**
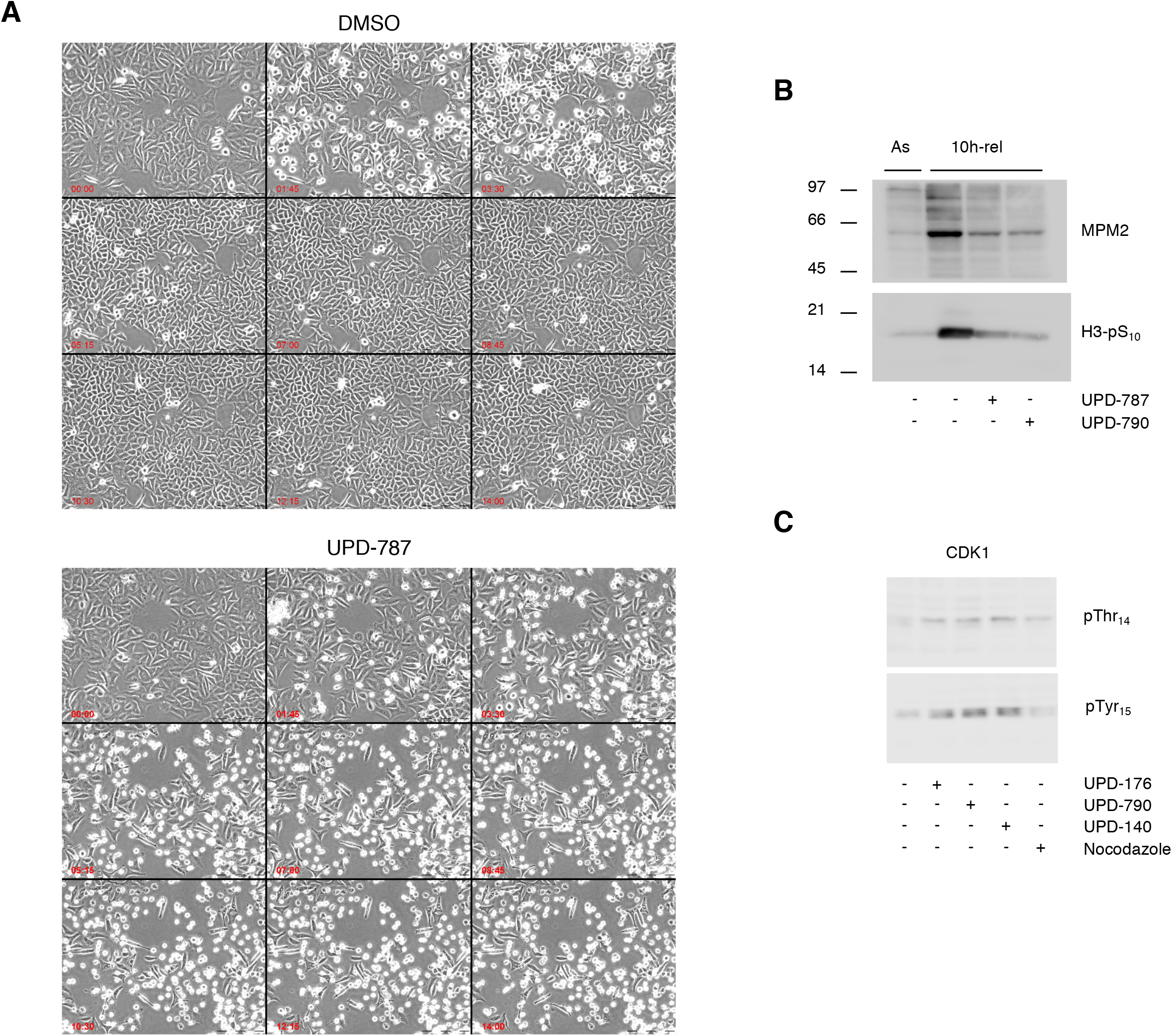
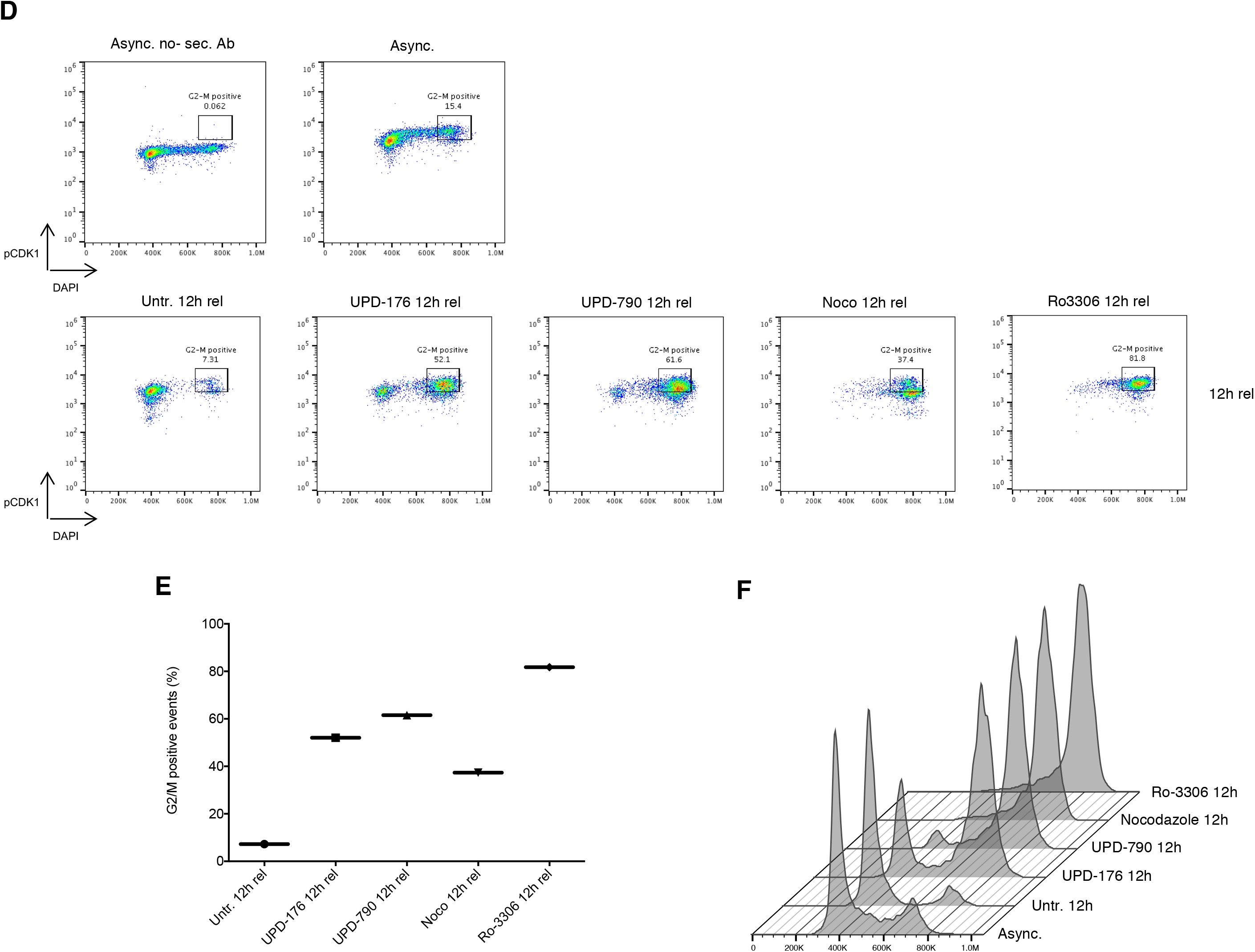
CDC25 inhibitors impair the execution of mitosis. (A) Phase contrast stills of HeLa cells synchronized by 2x thymidine block-release, treated with vehicle alone or UPD-787 (10 μM) 5h upon release and visualized from 7.5h to 21.5h upon release (4 frames/h). (B) Western blot analysis of MPM2 epitopes and Histone H3 phosphorylation (pSer_10_) in HeLa cells treated as in (A) and examined at 10h upon release from the 2x thymidine block. (C) Western blot analysis of CDK1 (pThr_14_ and Tyr_15_, respectively) from HeLa cells treated as in (A) and examined at 10h upon release from the 2x thymidine block. (D) Flow cytometric analysis of CDK1-pTyr_15_ in cells treated as in (A) and examined 12h upon release from the 2x thymidine block. Nocodazole (0.4 μg/ml) and Ro-3306 (9 μM) were used as controls for the level of CDK1 phosphorylation. Staining with secondary antibody only (top left panel) was used to subtract self-fluorescence. (E) Quantification of positive events gated as shown in (D). (F) DNA content (DAPI staining) of the cells shown in (d).

Biochemical analysis of mitotic markers at 10h from the double-thymidine block-release showed decreased MPM2 epitopes in treated cells, a read out for CDK1 activity, and reduced Histone H3 phosphorylation at Ser_10_, a read-out for chromosome condensation (Fig. 4B). To directly assess cellular CDK1 activity under these conditions we used antibodies recognizing phosphorylated Thr_14_ or Tyr_15_, two sites in the P-loop of CDK1 catalytic domain, the phosphorylation of which hampers kinase activity and that are selectively dephosphorylated by CDC25 (Atherton-Fessler, Parker et al., 1993, Ferrari, 2006). As expected, at 12h from the double-thymidine block-release point, control cells progressed to the G1 phase of the next cycle displaying low pThr_14_/pTyr_15_-CDK1, whereas cells treated with the compounds accumulated at G2/M (Fig. 4F) displaying high pThr_14_/pTyr_15_-CDK1 (Fig. 4C-E). The level of pTyr15-CDK1 in cells treated with CDC25 inhibitors appeared to be intermediate between that of cells treated with Ro-3306, a specific inhibitor of CDK1, and of cells treated with Nocodazole, an agent that interferes with tubulin polymerization causing arrest at pro-metaphase with active CDK1 (Fig. 4E).

### Effect of compounds on CDC25A overexpressing cell lines

To determine whether also cancer cells overexpressing CDC25 are sensitive to treatment with the compounds identified in this study, we focused on cell lines such as the lung carcinoma A549 that carry activating bi-allelic mutation of *K-Ras (G12S)* and are reported to express high level of CDC25B (http://www.proteinatlas.org/). However, Western blot analysis of cellular and nuclear extracts of A549, HeLa and Colo741 cells revealed lack of correspondence between the mRNA levels reported in databases and actual protein expression (Fig. S8). Similar results were obtained upon analysis of CDC25A and CDC25C protein expression (Fig. S8). Given the lack of an appropriate cell line model, we focused on a system that is characterized by tetracycline-dependent expression of CDC25A and has been previously described (Neelsen, Zanini et al., 2013). Viability assays showed that cells grown in the absence of tetracycline, a condition inducing ectopic expression of HA-CDC25A to high extent, remained sensitive to treatment with UPD-176 or UPD-787 (Fig. 5A and 5B).

**Figure 5.**
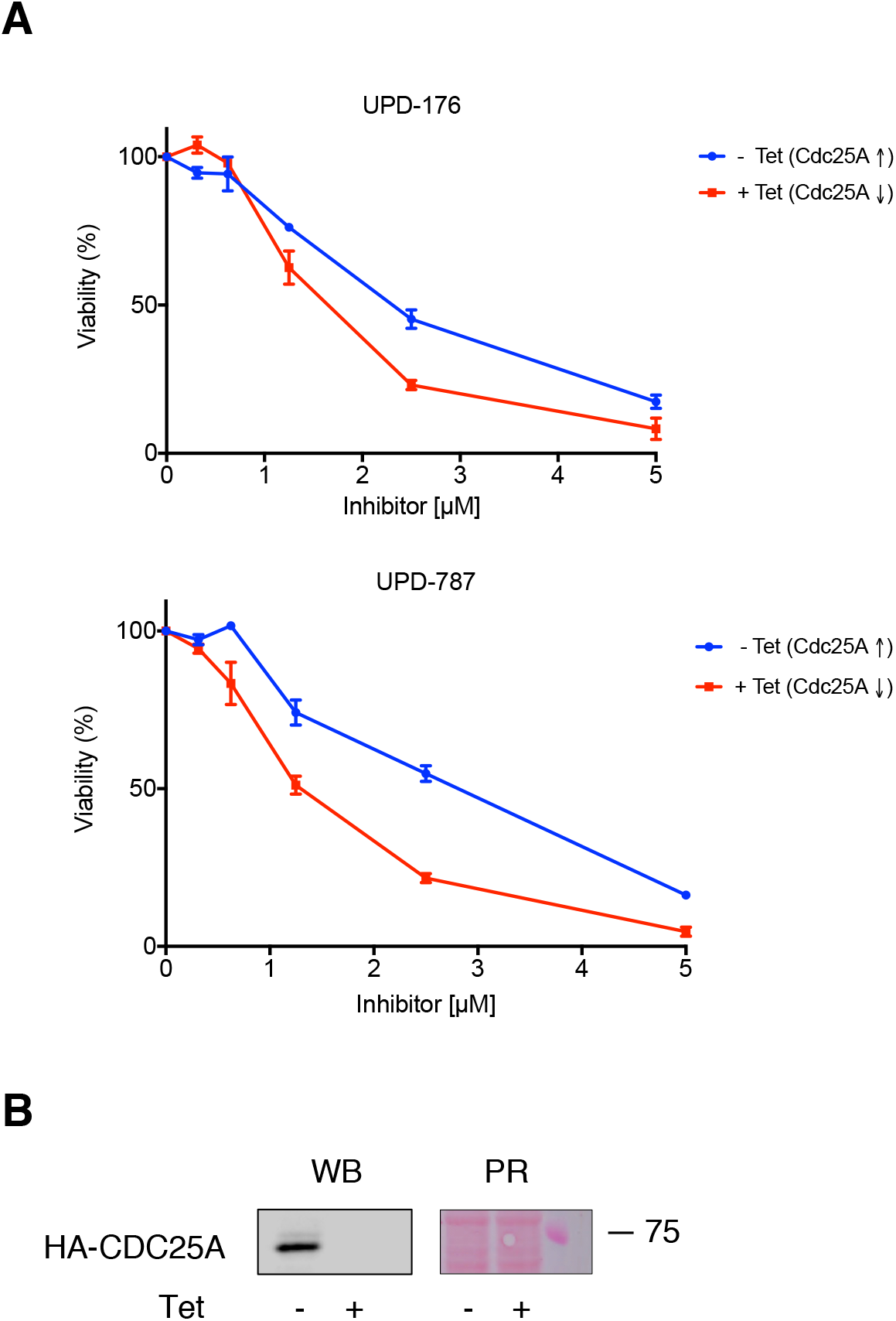
Cells overexpressing CDC25 are sensitive to CDC25 inhibitors. (A) U2OS Tet-OFF cells expressing HA-CDC25A were treated with the indicated compounds and cell viability was determined. (B) Western blot (WB) analysis of HA-CDC25A expression in the presence or absence of tetracycline. PR: Ponceau Red staining.

### Effect of CDC25 inhibitors on the growth of *Apc/K-Ras* intestinal organoids

Having established the effect of CDC25 inhibitors on the growth of cells in monolayer cultures, we addressed the ability of selected compounds to affect the growth of organoid cultures. Since CDC25A cooperates with, and is rate-limiting in tumorigenesis induced by *Ras* (Ray & Kiyokawa, 2008), we generated and cultured 3D-organoids from stem cells of the intestinal crypts of *Apc/K-Ras* mice (Moor, Anderle et al., 2015). The latter carry the loss-of-function mutation *Apc1638N* combined with the villinCre-driven gain-of-function *KrasG12V* mutation (Valenta, Degirmenci et al., 2016). Upon seeding *Apc/K-Ras* organoids along with UPD-176 (5 μM), we observed that growing spheroids partially invaginated (Fig. 6A), indicating reduction of stem cell renewal accompanied by increased differentiation (Grabinger, Luks et al., 2014), whereas higher doses of the compound caused massive death (data not shown). Consistent with the pattern of invagination at low compound treatment, immunofluorescence confocal imaging revealed augmented expression of lysozyme, a marker for cells exiting the proliferative compartment of the crypt and acquiring a differentiated state (Fig. 6B). Quantitative RT-PCR performed on the stemness marker Lgr5 and on the differentiation markers Lysozyme and Cryptdin confirmed the confocal microscopy data (Fig. 6C).

**Figure 6.**
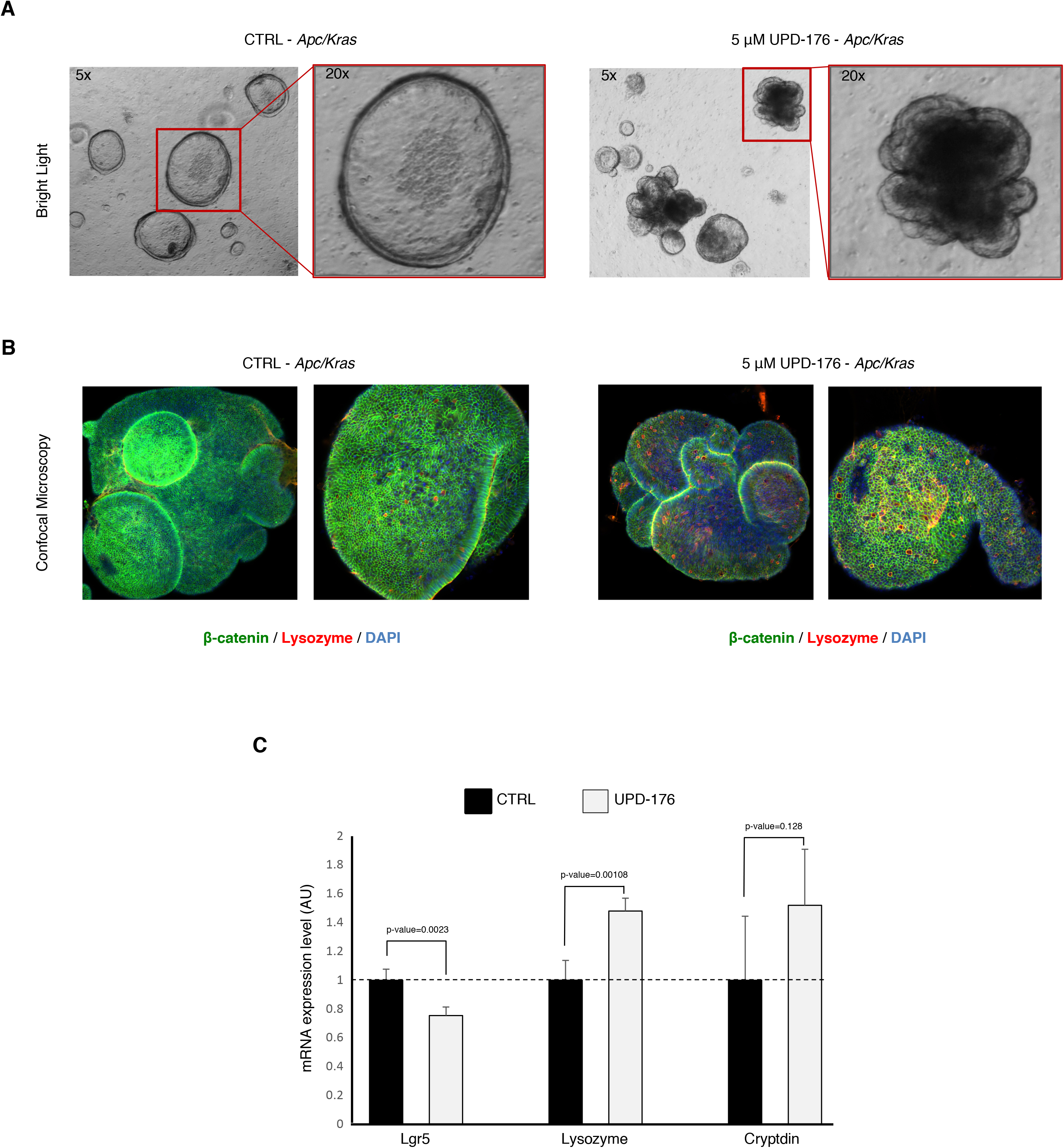
CDC25 inhibitors block the growth of K-Ras-dependent intestinal organoids. (A) Organoids reproducing duodenal crypts and derived from *Apc/Kras* mouse small intestine were treated with the UPD-176 (5 μM). Phase contrast stills were obtained at 48h of treatment. (B) Confocal immunofluorescence microscopy of *Apc/Kras* organoids treated as in A and stained for proliferation (β-catenin) or differentiation (lysozyme) markers. DNA was stained with DAPI. (C) qRT-PCR on cDNA obtained from control- and UPD-176-treated organoids was conducted using primers to the indicated markers.

## DISCUSSION

Precision oncology is centered on the principle of continuous molecular interrogation of tumors and on the use of dedicated pharmacological tools to maximize success in therapy. Constant monitoring is essentially intended to assess resistance (Komarova & Wodarz, 2005) and reveal pathways to which tumors may become addicted during treatment (Luo, Solimini et al., 2009).

CDC25 phosphatases are overexpressed in a variety of human cancers (Boutros et al., 2007, Kristjansdottir & Rudolph, 2004), are rate-limiting in tumorigenesis induced by *Ras* (Ray & Kiyokawa, 2008), and were recently proposed as target of choice in unresponsive, triple-negative breast cancer (Liu et al., 2018). In this study we set out to identify novel CDC25 inhibitors. Taking advantage of knowledge gained in previous drug discovery programs (Lavecchia, Di Giovanni et al., 2012a, Pestell et al., 2000), we initially defined a pharmacophore that is common to compounds belonging to three distinct classes of established CDC25 inhibitors (Fig. S1). Performing a ligand-based virtual screen of a proprietary library, we identified novel naphthoquinone inhibitors of CDC25 phosphatases (Fig. 1, Tables S1 and S2). Enzymatic studies revealed a mixed mechanism of inhibition for all CDC25 members (Fig. S2), possibly indicating binding of the compounds to catalytic (Fig. 1C) and allosteric sites of the phosphatase, as also reported for other CDC25 inhibitors (Bana et al., 2015, Brault, Denance et al., 2007, Lavecchia et al., 2012b, Lazo et al., 2001, Lazo, Nemoto et al., 2002). Mechanistically, we observed that compounds acted reversibly on CDC25 (Fig. 2C), likely through oxidation of the catalytic cysteine (Fig. 2B), a mode of action that we confirmed in cells (Fig. 3B). Such mechanism is compatible with a model proposed for CDC25 inhibitors (Sohn & Rudolph, 2003), according to which the thiolate group of the active-site cysteine undergoes very rapid conversion to sulfenic acid and it is protected from further conversion into irreversibly inactivated sulfinic acid by a back-door cysteine located in close proximity to the catalytic residue.

Cellular experiments conducted in non-synchronous or synchronized cells showed that the most potent compounds arrested cells at the G1/S or the G2/M transition (Fig. 3D and 3E). In cells synchronized with double-thymidine treatment, if compounds were administered at the time of release from the block (Fig. S5B), a minimal progression to S-phase appeared evident upon 6 hours of release. This may indicate that either turnover of the compound occurred during this time or, more likely, that the amount of CyclinE/CDK2 complex built up while cells accumulated at the point of forced arrest (i.e., early S-phase) was sufficient to support cell cycle progression despite inhibition of CDC25. The latter hypothesis is corroborated by the observation that when compounds were added in mid-G1, transition to S-phase was effectively blocked (Fig. 3D).

Since quinones undergo reduction to form semiquinone radicals or hydroquinones that, in turn, can react with oxygen to form superoxide radicals and/or hydrogen peroxide (Brezak, Kasprzyk et al., 2008), hence affecting a number of metabolic processes in the cell, we examined DNA damage induction as a possible side-effect relevant to the overall cellular response observed. Specifically, we quantified the biomarker γ-H2AX in synchronized cells that were treated with compounds. The data excluded the possibility that the cell cycle arrest observed in response to compound treatment could be secondary to genotoxic effects (Fig. S6).

To examine the cellular mechanism of action for the compounds described in this study, we focused on activation of CDK1 and entry into mitosis. Analysis of the extent of Thr14/Tyr15 phosphorylation, as indicator of kinase activation, and phosphorylation of CDK1 substrates, as indicator of kinase activity, revealed that the inhibitors effectively impaired both responses (Fig. 4B-E). Chromosome condensation normally occurring in prophase was also impaired in compound-treated cells, in line with flow cytometry data attesting a G2/M arrest under these conditions (Fig. 4B and 4F). Visual inspection of cells treated with the compounds showed that they could not execute mitosis, in line with our biochemical evidence, but rather underwent massive death (Fig. 4A). Administration of low concentration of the inhibitors to HeLa cells carrying mCherry-H2B allowed appreciating failed attempts to round up for mitosis, followed by membrane blebbing and death (Fig. S7, movies M2 and M3).

Enzymatic profiling of two of the most potent CDC25 inhibitors on a panel of phosphatases revealed PP5 as the only other target of the compounds (Table S4). PP5 controls a number of cellular processes including proliferation, migration and DNA damage. Interestingly however, PP5 activity is normally off due to folding of an N-terminal inhibitory domain onto the catalytic site, with ligand-mediated release of auto-inhibition occurring in response to cellular cues (Shi, 2009). Hence, we argue that the specific G1/S and G2/M responses that we describe in this study are genuine effects of the identified compounds on CDC25 phosphatases.

Considering that CDC25A and CDC25B are overexpressed in a variety of human cancers (Boutros et al., 2007) and *CDC25A* was demonstrated to be a rate-limiting oncogene in transformation by *RAS* (Ray & Kiyokawa, 2008), we examined the response of a cell line overexpressing CDC25A, as paradigm for the demonstration of compounds potency. The data confirmed that the compounds could effectively decrease viability of CDC25 overexpressing cells (Fig. 5).

Finally, we conducted studies in 3D-organoids, a system that reproduces architecture and function of the tissue of origin in reduced scale (Kingwell, 2016). Organoids obtained from stem cells of the intestinal crypts of *Apc/K-Ras* mice feature an internal cavity, corresponding to the crypt’s lumen, but lack the symmetry characteristic of this organ (Sato, Vries et al., 2009). Fluorescence microscopy revealed decreased organoids size and acquisition of a differentiated state in response to CDC25 inhibitors, a pattern confirmed by qRT-PCR on selected markers (Fig. 6). These data are reminiscent of observations made on *Cdc25B^−/−^/Cdc25C^−/−^* knock-out mice where *Cdc25A* was conditionally disrupted (*Cdc25A^fl−^*) in all tissues of the adult organism (Lee et al., 2009). In this triple knock-out background, the authors reported large loss of the small intestine and crypts atrophy due to arrest at G1 and G2 phases of the cell cycle, which was paralleled by an increase of epithelial cell differentiation. As a whole, the triple knock-out studies and our data support the concept that blocking cell cycle progression through inhibition of CDC25 activity is beneficial to target tumor growth driven by mutant *Ras* and dependent on CDC25. Hence, considering the failure of previous attempts, it is desirable to revitalize drug discovery programs on CDC25 inhibitors. The studies on naphthoquinone scaffolds described here, along with a follow-up program that we are conducting on *ad hoc* synthesis of derivatives, open interesting perspectives to the design of novel anti-cancer therapeutics targeting CDC25.

## MATERIALS AND METHODS

### Microscopy

Cells were grown in 35 mm CellView™ cell culture dishes with glass bottom (# 627870, Greiner-BioOne) at a density of 2.5 × 10^5^ cells/ml in a humidified cell incubator maintained at 37 C° and 5% CO_2_. Cell were viewed by phase contrast microscopy with a 10x objective using an Olympus IX 81 motorized inverted microscope (Olympus, Hamburg, Germany) equipped with external temperature control chamber and CO_2_ bottle to maintain cells at 37°C with 5% CO_2_. Transition through mitosis was documented by acquisition of four frames per hour over a period of 14h using a CCD camera (Orca AG, Hamamatsu) and cellR^®^ software (Olympus). HeLa-Kyoto cells (mCherry-H2B/EGFP-a-tubulin) were visualized by fluorescence microscopy with an Olympus IX 81 microscope using a 10x objective and selecting Ex. = 492 ±18 nm / Em. = 535 ±50 nm for the green channel and Ex. = 572 ±23 nm / Em. = 645 ±75 nm for the red channel.

### Intestinal organoids culture

Intestinal crypts were isolated from *Apc/Kras* animals bearing *Apc^1638N^* and villin-driven *Kras^G12V^* alleles with slight modifications (Valenta et al., 2016) of a previously described method (Sato, Stange et al., 2011). Briefly, crypts were isolated and purified, embedded in 50 μl matrigel drops (Corning, 356231) and overlaid with 500 μl organoid medium Advanced DMEM/F12 (Life Technologies, 11320-082) containing 2 mM GlutaMAX (Life Technologies, 35050-061), 10 mM HEPES buffer (Sigma, 83264-100ML-F), 0.5 U/ml Penicillin/Streptomycin (Life Technologies, 15070-063), N2 (Life Technologies, 17502-048), B27 (Life Technologies, 12587-010)]. Organoids were expanded for 5 days in growth factor supplemented medium containing 50 ng/ml mEGF (Life Technologies, PMG8041), 100 ng/ml mNoggin (Peprotech, 250–38) and 500 ng/ml hRSPO1 (R&D, 4645-RS-025). EGF and R-spondin were subsequently removed to select for organoids that had lost the wild-type *Apc* copy. The medium was changed every 2 days. Established mutant lines were passaged every four days by mechanical disruption with a bent P1000 pipette tip.

## ACKNOWLEDGMENTS

We would like to thank: I. Hoffmann (DKFZ, Heidelberg, Germany) for CDC25A expression constructs; B. Gabrielli (University of Queensland, Australia) for CDC25B and CDC25C constructs; N. J. Lamb (University of Montpellier, France) for suggestions on expression, and purification of CDC25 phosphatases; D. Gerlich (Austrian Academy of Sciences, Vienna, Austria) for Kyoto HeLa cells; the Molecular Modeling Section (MMS) of the University of Padua for kind support; F. M’hmedi for technical support and J. Nüssel for help with cellular assays.

This work was supported by grants from: Promedica-Stiftung-UBS, Stiftung für Krebsbekämpfung and Stiftung für wissenschaftliche Forschung of the University of Zurich to SF; Swiss National Science Foundation to KB; Forschungskredit of the University of Zurich to CC; AIRC (IG 14180) to LAP; Progetto Giovani Ricercatori, University of Padua, to GC.

## AUTHOR CONTRIBUTIONS

GR, GZ: designed and synthesized all compounds;

GC: performed molecular modeling and docking studies;

ZK, SK, CK, CG: performed experiments in Fig. 2-3-4-5, S2-S3-S7-S8 and movies M1-M3;

CC, JT: performed experiments in Fig. 6;

KB, LAP: contributed to the conceptual development of the study;

SF: wrote the manuscript.

